# Structure-guided antisense oligonucleotides selectively modulate frameshifting of a human gene

**DOI:** 10.64898/2026.09.03.748613

**Authors:** Ondrej Kostov, Myriam Moreno Swanton, Katie R. Waldon, Autumn M. Matthews, Marija Ciba, Mathias B. Danielsen, Balazs Schafer, Saheli Ganguly, Christopher C. Ebmeier, Marvin H. Caruthers, Alexandra M. Whiteley

## Abstract

Programmed -1 ribosomal frameshifting (-1 PRF) is a conserved translational recoding mechanism that expands proteomic diversity and regulates gene expression through RNA structural elements, most notably stimulatory pseudoknots. This mechanism is common in viruses, where it is used to control stoichiometry of viral protein products generated by the host cell to direct viral replication. Despite its biological importance, strategies to selectively modulate frameshifting remain limited. The mammalian retrotransposon-derived gene *PEG10* also relies on -1 PRF to produce a fusion protein, gag-pol, which is necessary for reproduction but has also been implicated in neurological diseases. Here, we establish an antisense oligonucleotide (ASO) targeting an RNA structural element as an effective approach to tune *PEG10* frameshifting. Using structure prediction, systematic antisense tiling across the *PEG10* pseudoknot, and multiple model systems, we identify a discrete functional hotspot within the lower RNA stem that governs frameshift efficiency. ASOs targeting this region selectively suppress gag-pol production with minimal impact on gag, thereby shifting the ratio of protein products in a dose-dependent manner. Mechanistic dissection using RNase H-active and -inactive ASO designs, pre-annealed duplexes, and fluorescence-based subcellular localization supports a predominantly nuclear mode of action in which ASOs engage nascent *PEG10* transcripts and bias pseudoknot folding away from the frameshift-competent conformation. Functional effects are conserved between human cell lines and murine models, including neurons, highlighting the generality of this strategy. Together, our results define RNA structural dynamics as a druggable layer of translational regulation and establish antisense modulation of pseudoknot folding as a way to control endogenous frameshifting. This work provides a conceptual and practical framework for targeting recoding-dependent gene products such as PEG10 in disease and suggests broader applicability of structure-directed ASOs to viral and cellular frameshifting elements.

## Introduction

Antisense oligonucleotides (ASOs) have emerged as a powerful therapeutic modality for precise modulation of gene expression, enabling sequence-specific targeting of disease-associated transcripts without the need for permanent genomic alteration (Bennett 2019). Clinical successes, including splice-modulating ASOs for spinal muscular atrophy (SMA) (Singh et al. 2006; Corey 2017; Passini et al. 2011) and knockdown ASOs for *SOD1*-associated amyotrophic lateral sclerosis (ALS) (Benatar et al. 2025; Miller et al. 2022), have validated the therapeutic potential of oligonucleotide-based drugs (Lauffer et al. 2024).

Most approved ASO strategies rely on transcript degradation or splicing modulation and are not well suited for therapeutic scenarios in which selective tuning of protein translation, rather than total gene silencing, is required. This is particularly relevant for viruses, which often utilize multiple reading frames of the same mRNA to generate distinct protein products necessary for different stages of the viral lifecycle (Firth and Brierley 2012). This is accomplished through programmed ribosomal frameshifting (PRF), where ribosomes stall on specific RNA sequences, either defined by structural elements (Brierley et al. 2007) or by viral and host protein factors (Napthine et al. 2017, 2021), and induce a change in the reading frame. The frequency of PRF is unique to individual viruses and fine-tuned to promote their replication, and perturbation of frameshifting ratios often results in profound replication defects (Karacostas et al. 1993; Park and Morrow 1991; Sun et al. 2021). With this in mind, there has been considerable interest in identifying pharmaceutical candidates that can modulate PRF as a therapeutic strategy for a variety of human viruses (Kelly et al. 2021).

Structured RNA elements, including pseudoknots, can exhibit conformational plasticity and dynamic remodeling during translation that are important determinants of frameshift efficiency (Giedroc and Cornish, 2009). Biophysical and single-molecule studies have shown that frameshifting efficiency correlates more closely with the ability of pseudoknots to sample alternative conformations than with their simple mechanical resistance to ribosomal unfolding (Ritchie et al., 2012). Antisense oligonucleotides therefore provide a potential means of modulating frameshifting by perturbing RNA secondary and tertiary structure, either by creating alternative ribosome-stalling structures or by sequestering base-pairing interactions required for formation of the native pseudoknot. Although antisense-mediated modulation of ribosomal frameshifting has been demonstrated in viral and reporter systems (Kibe et al. 2025; Zhang et al. 2021; Kelly et al. 2021; Howard et al. 2004; Henderson et al. 2006), the selective suppression of endogenous frameshifting in a mammalian gene remains comparatively unexplored.

About 40% of the human genome consists of transposable elements, many of which also utilize PRF to generate multiple protein products from the same mRNA (International Human Genome Sequencing Consortium et al. 2001; Lawson et al. 2023; Atkins et al. 2016). In rare cases throughout evolutionary history, these elements have been domesticated, or co-opted, by the host to perform adaptive functions (Volff 2006; Jangam et al. 2017;. Frank and Feschotte 2017). PEG10 is an imprinted mammalian gene (Ono et al. 2001) derived from a domesticated retrotransposon (Brandt et al. 2005b, 2005a) and is essential for placental development (Ono et al. 2006; Shiura et al. 2021). Like viruses, PEG10 utilizes programmed −1 ribosomal frameshifting (−1 PRF) to generate two protein isoforms from a single mRNA: a shorter gag-like protein (RF1) and a longer gag–pol fusion protein (RF1/2) (Clark et al. 2007; Lux et al. 2010). Frameshifting is directed by a highly conserved slippery heptamer sequence (GGGAAAC) followed by a downstream RNA pseudoknot (Manktelow 2005), yielding a defined gag-pol:gag production ratio in mammalian cells. The strict conservation of PEG10 frameshifting across mammals (Brandt et al. 2005b; Black et al. 2023) underscores its functional importance: recent studies have revealed that dysregulation of PEG10 frameshifting in either direction is maladaptive. First, functional mutation of the pol region’s protease domain is embryonic lethal in mice due to defects in placental development (Shiura et al. 2021). Similarly, forced expression of PEG10 gag-pol in the absence of gag is also developmentally lethal (Shiura et al. 2025). In summary, careful balance of PEG10 translation is essential for biological function.

Dysregulation of gag-pol:gag abundance is also evident in human disease. In Angelman syndrome, loss of functional UBE3A leads to elevated levels of gag-pol protein, thereby altering neuronal trafficking in embryonic brains (Pandya et al. 2021). In ALS, dysfunction of the proteasomal shuttle factor UBQLN2 also selectively impairs degradation of PEG10 gag-pol (Whiteley et al. 2021; Black et al. 2023; Roberts et al. 2025). Gag-pol contains a protease and undergoes protease-dependent self-cleavage (Golda et al. 2020; Clark et al. 2007; Lux et al. 2010), releasing a small protein fragment that translocates to the nucleus and alters transcriptional programs involved in neuronal maintenance and axon remodeling (Black et al. 2023). Together, these findings position PEG10 frameshifting, not total PEG10 expression, as a therapeutically actionable control point.

Conventional oligonucleotide strategies are poorly suited for this target, and small molecules that target frameshifting in other genetic elements have little or no effect on *PEG10* (Cardno et al. 2015; Sun et al. 2021). Gapmer ASOs and siRNAs reduce total *PEG10* mRNA levels and thereby suppress both gag and gag–pol, risking disruption of essential PEG10 functions. Splice-switching ASOs are inapplicable, as the *PEG10* frameshift occurs within a continuous open reading frame rather than at exon–intron junctions. These limitations motivate a fundamentally different approach: direct modulation of the *PEG10* frameshift signal itself to selectively rebalance gag and gag-pol production. In support of this strategy, ASOs can be used to both positively and negatively influence the frameshifting rate of HIV (Vickers and Ecker 1992).

Here, we present a structure-guided antisense strategy to selectively inhibit *PEG10* −1 PRF and suppress production of the disease-associated gag-pol isoform while preserving basal gag expression. By systematically mapping antisense oligonucleotide placement across the *PEG10* pseudoknot and comparing multiple ASO chemistries, we identify non-degradative oligonucleotides that selectively modulate frameshifting without inducing transcript degradation. We further provide evidence for nuclear localization of these ASOs and a mechanism consistent with their engagement with *PEG10* RNA during an early stage of RNA biogenesis. Together, these results establish programmable RNA structure modulation as a viable therapeutic strategy for selectively rebalancing PEG10 protein isoforms and suggest a general framework for targeting structured regulatory elements within endogenous human mRNAs.

## Results

### The *PEG10* mRNA pseudoknot is amenable to antisense oligonucleotide targeting

To enable rational antisense targeting of the *PEG10* frameshift signal, we first established a structural framework for the *PEG10* pseudoknot and its dynamic formation. The *PEG10* −1 PRF element consists of a conserved slippery heptamer followed by a compact H-type pseudoknot whose architecture is required for efficient frameshifting and production of two distinct proteins (**Figure 1a**). Published chemical probing and mutational analyses (Manktelow 2005) have defined the core stem–loop topology of this structure, while recent structure prediction methods with Alphafold 3.0 (Abramson et al. 2024) suggest a compact tertiary fold with closely apposed stem–loop junctions (**Figure 1b**).

**Figure 1:**
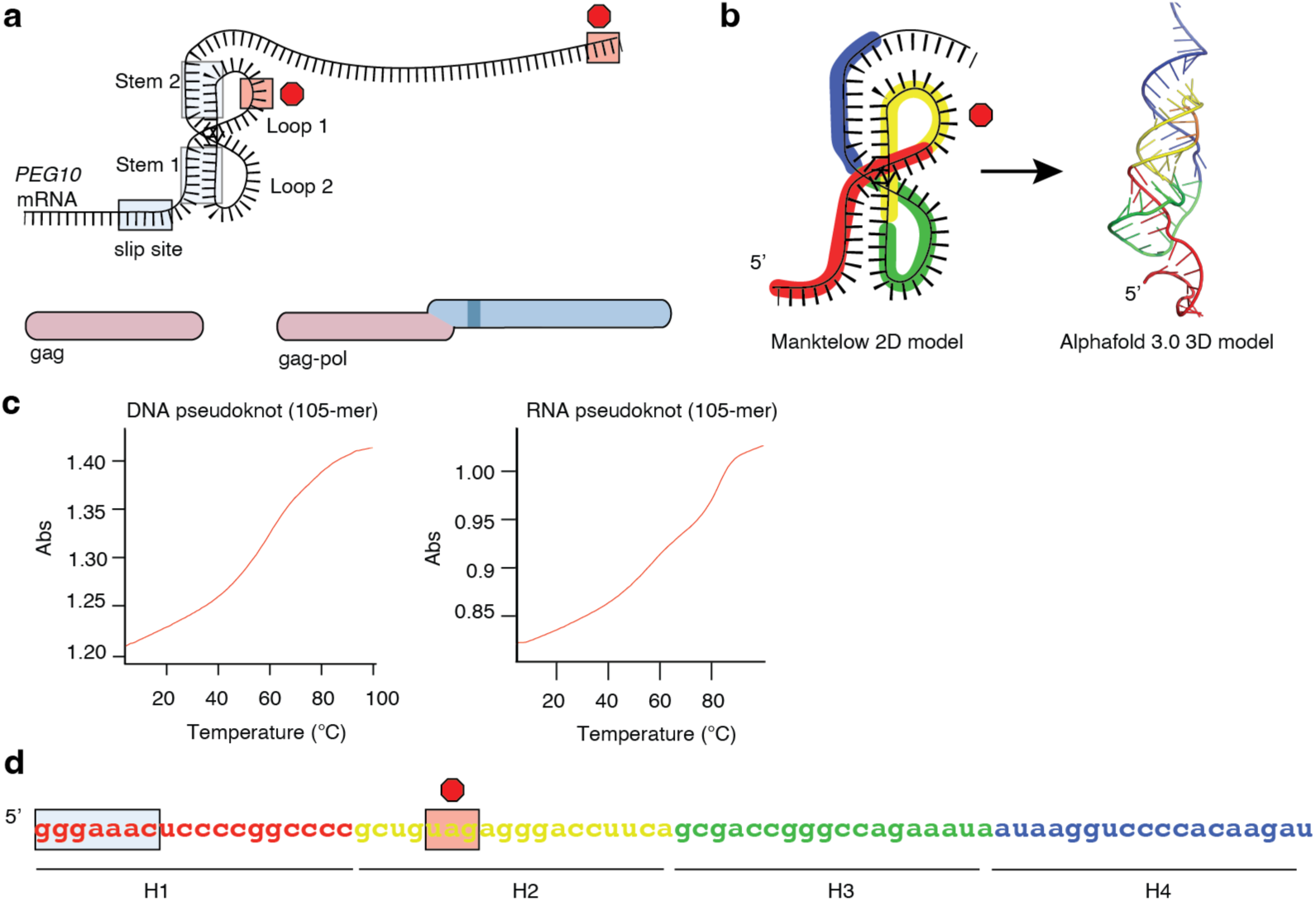
The *PEG10* mRNA pseudoknot is amenable to antisense oligonucleotide-based modulation. **a)** Schematic of published structure of *PEG10* mRNA pseudoknot with resulting protein products. The stop codon at the end of gag is found in Loop 1, and the later stop codon at the end of gag-pol is shown at the far right (not to scale). If the first stop codon is used, gag protein is created; if the ribosome slips -1 due to the pseudoknot, the second stop codon is used to generate gag-pol fusion protein. The dark blue region of pol denotes the protease domain found only in gag-pol protein. **b)** Left: color-coded schematic of the Manktelow et al. preferred model of PEG10 pseudoknot. Right: Alphafold 3.0 3D model of *PEG10* pseudoknot structure. Relevant loops and Stems from (a) are highlighted. **c)** thermal denaturation profiles of the 105-nt synthetic pseudoknot sequence prepared as DNA (left) or RNA (right), demonstrating distinct folding behavior and melting transitions consistent with chemistry-dependent secondary structure formation. **d)** Sequence of *PEG10* pseudoknot RNA with regions of TMO sequence complementarity shown. These complementary sequence regions are highlighted by H1, H2, H3, and H4. The slippery sequence is highlighted in blue box, and the stop codon of the gag ORF is highlighted in red. Sequence colors match those in (b).

To experimentally model the *PEG10* frameshift structure, we designed a synthetic 105-nt RNA construct encompassing the full pseudoknot region and flanking sequences, as well as corresponding DNA analogs for biophysical characterization (**Figure 1c**). Both RNA and DNA constructs were prepared in high purity, as confirmed by analytical HPLC and mass spectrometry (**Supplementary Figure 1**). Thermal denaturation analyses and structure prediction revealed distinct folding behaviors between RNA and DNA versions of the construct, consistent with differential secondary structure formation and highlighting the sensitivity of the pseudoknot to subtle perturbations in backbone chemistry and base pairing.

We next performed an antisense “tiling” screen across the *PEG10* pseudoknot to identify regions most amenable to functional modulation of frameshifting. ASOs were designed to target discrete structural elements of the pseudoknot, including sequences proximal to the slippery site, stem regions, and loops (**Figure 1d**).

### RNase-insensitive ASOs alter frameshifting of *PEG10* in cells

ASOs were then administered to cells stably expressing a *PEG10* frameshift reporter. To begin, thiomorpholino ASOs (TMOs) were chosen because they do not activate RNase H or RNAi pathways (Le et al. 2022; Langner et al. 2020) and therefore would not lead to degradation of the *PEG10* mRNA. Flp-in HEK293 cells were stably transfected with a PEG10 reporter containing a fusion with mCherry at the N-terminus of gag and a fusion with eGFP at the C-terminus of gag-pol (**Figure 2a**). Cells expressing PEG10 make a mixture of gag and gag-pol products dependent on PRF, with gag represented by red fluorescence, and the latter frameshifted product exhibiting both red and green fluorescence. After selection for stable integrants, these cells were transfected with 50 nM ASOs directed against different segments of the *PEG10* pseudoknot. 72 hours later, cells were examined by microscopy. Cells transfected with H1 and H2 TMOs showed no major difference in fluorescence compared to control, whereas H3 and H4 TMO-transfected cells showed a marked shift towards predominantly red fluorescence (**Figure 2b**).

**Figure 2:**
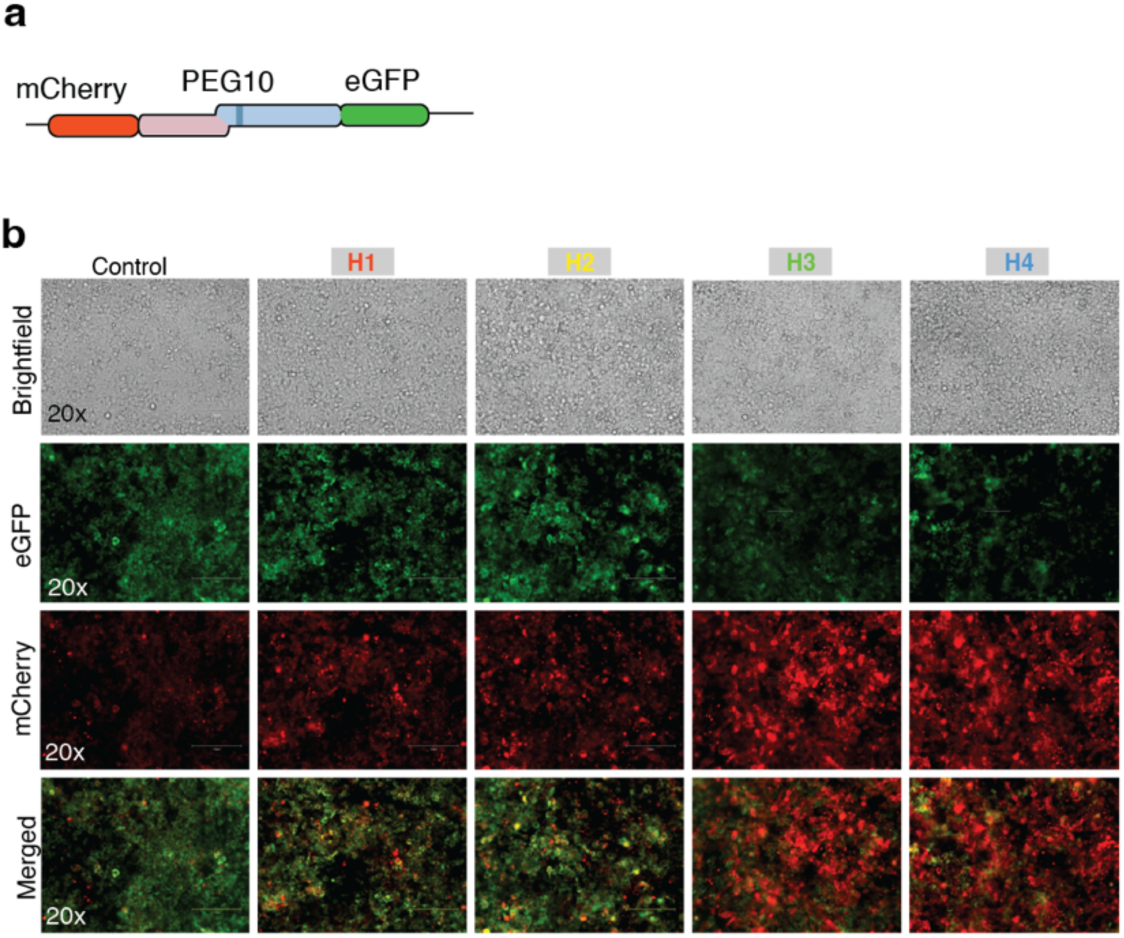
Antisense oligonucleotides efficiently prohibit *PEG10* frameshifting in cells. **a)** Schematic of *PEG10* frameshift reporter. PEG10 gag is fused at the 5’ end to mCherry, and at the 3’ end to eGFP. Translation of gag results in red fluorescence, and translation of gag-pol results in red and green fluorescence. **b)** Microscopy of cells stably transfected with frameshifting reporter and acutely transfected with *PEG10* ASOs. Cells were imaged 72 hours after transfection with 50 nM ASO in Lipofectamine 2000. Images were collected with the same light intensity and gain for each well. Images are from two representative experiments.

The same TMOs were then tested for their ability to regulate frameshifting of the endogenous *PEG10* mRNA in HEK293 cells. PEG10 gag-pol protein is rapidly degraded by the proteasome dependent on the shuttle factor UBQLN2 (Black et al. 2023; Roberts et al. 2025), which limits its visibility by western blot. Therefore, HEK293 cells lacking expression of Ubiquilin 1, Ubiquilin 2, and Ubiquilin 4 (Itakura et al. 2016) were used in order to better visualize endogenous gag-pol protein. 72 hours after transfection using Lipofectamine 2000, H3- and H4-treated cells showed a dose-dependent effect on gag-pol expression with minimal effects on gag (**Figure 3a-c**), which was most clearly visualized by directly plotting the ratio of gag-pol:gag protein per condition (**Figure 3d**). Therefore, TMO treatment influences PEG10 protein production from endogenous *PEG10* mRNA as well as the engineered reporter construct. Based on these data, we conclude that TMOs targeting pseudoknot formation that hybridize with sequences after the gag stop codon (H3 and H4) are potent inhibitors of PRF.

**Figure 3:**
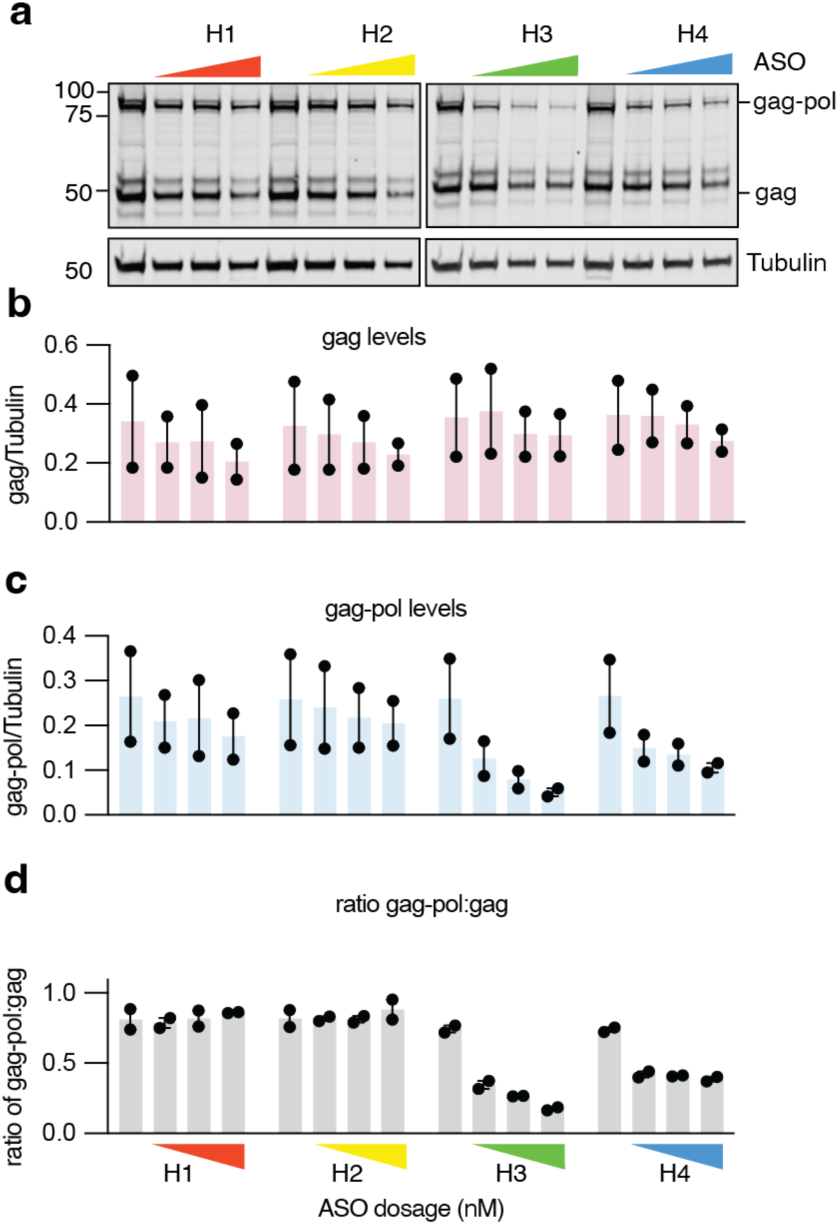
Antisense TMOs efficiently prohibit endogenous *PEG10* frameshifting. **a)** Western blot showing the effects of increasing doses of each ASO sequence on endogenous PEG10 gag and gag-pol expression of HEK293 cells. Cells were transfected with 50, 100, or 200 nM of each ASO using Lipofectamine 2000 and harvested 72 hours later. Shown is one representative of two blots. **b-c)** Quantification of gag (b), and gag-pol (c). **d)** Ratio of gag-pol:gag from two independent experiments. Data are from two representative experiments.

To further refine the positional requirements within this sensitive region, we designed extended variants of the H3 TMO that encroached either toward the apical hairpin region (H2) or toward the downstream second stem (H4). Extension of the H3 binding site in either direction did not meaningfully influence the TMO’s ability to influence the gag-pol:gag ratio (**Supplementary Figure 2**), indicating that maximal functional efficacy involves positioning of the TMO somewhere within the lower stem region. These data highlight a narrow “hot spot” within the pseudoknot where antisense engagement is optimally coupled to functional modulation of frameshifting.

### Multiple chemistries support inhibition of PEG10 frameshifting

The chemistry of antisense oligonucleotides has radiated to include modification of sugar moieties, modification of the phosphate backbone, and mixed chemistries that enhance bioavailability, toxicity, and potency (Bennett 2019). To compare the ability of different ASO chemistries to inhibit frameshifting, cells were transfected with a variety of ASO types (**Figure 4a**) using the same H3 sequence. As a negative control, the ‘sense’ sequence of H3 in RNA format was transfected into cells. Transfection of all tested chemistries had no effect on levels of gag protein (**Figure 4b-c**). In contrast, gag-pol levels were profoundly decreased by TMO transfection, and were also decreased by PS-MOE and PS-LNA transfection (**Figure 4b,d**). Upon comparison of the gag-pol:gag ratio of PEG10 across conditions, strong inhibition of frameshifting was observed with MOE and PS-LNA in addition to TMOs (**Figure 4e**). To complement these findings, the different modified ASOs were hybridized to *PEG10* RNA and the thermal stability of duplexes was measured. The duplex with phosphorothioate RNA ASO had a melting temperature of 83.0°C (**Figure 4f**). In comparison, TMOs had the closest melting temperature at 82.2°C while PS-OMe, PS-MOE, and especially PS-LNA exhibited elevated melting temperatures (**Figure 4f**).

**Figure 4:**
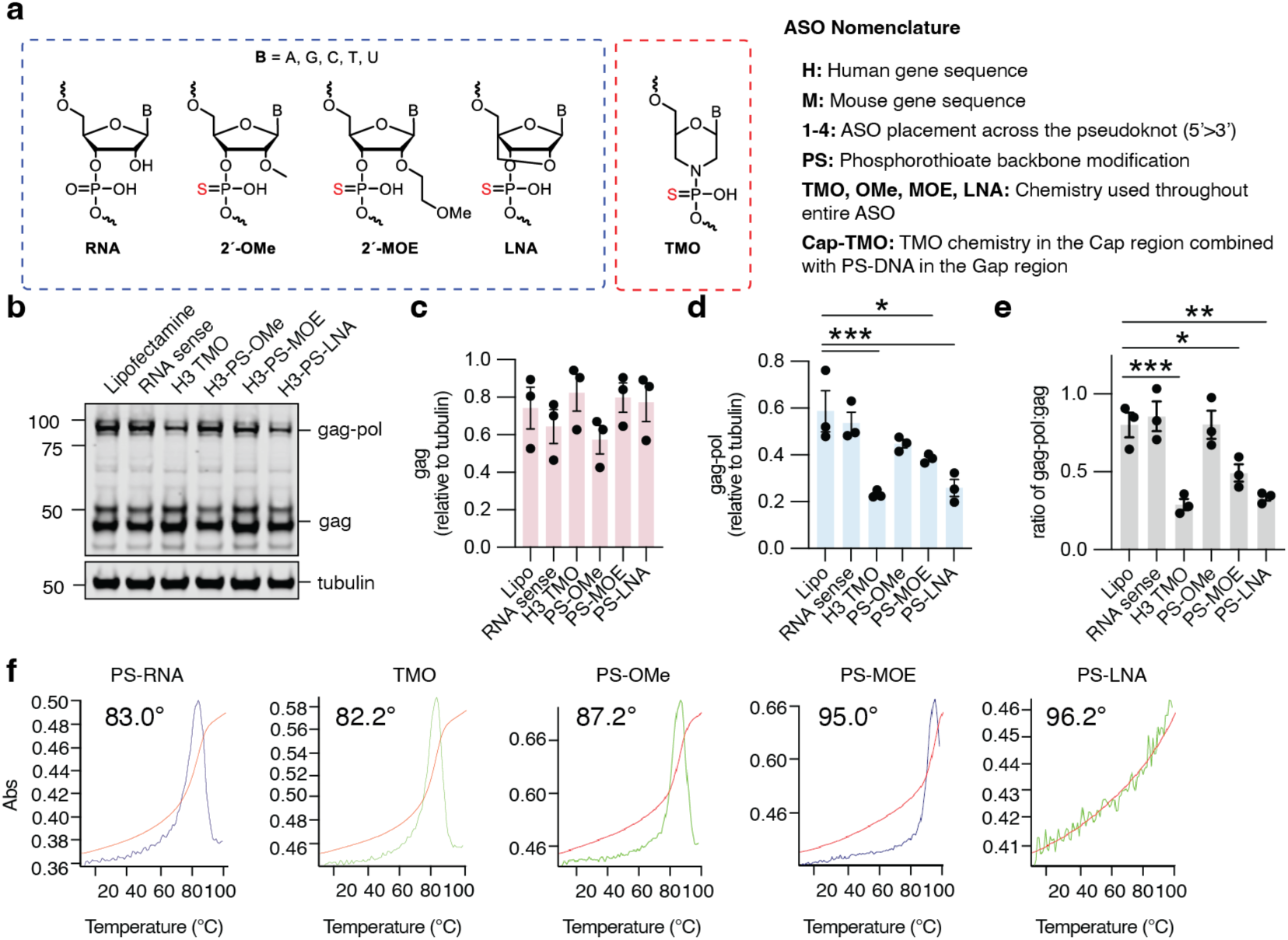
Alternative H3 chemistries result in different mRNA fates. **a)** Schematic of different ASO chemistries tested, with ASO nomenclature detailed on the right. **b)** Representative western blot of HEK293 cells transfected with each H3 chemistry for levels of PEG10 gag-pol and gag. Shown is one of 3 representative blots. **c-d)** Quantification of gag (c), and gag-pol (d). **e)** Ratio of gag-pol:gag from 3 independent experiments. For (c-e), statistics were determined by one-way ANOVA with multiple comparisons. **f)** Thermal melting temperatures (Tm) of each ASO hybridized to the complementary *PEG10* RNA sequence, illustrating chemistry-dependent differences in duplex stability. Images are representative of 3 experiments.

### Subcellular localization of ASO–mRNA interactions supports a nuclear site of action

We next examined the intracellular site of action of the leading H3 ASO candidate. In principle, ASOs could modulate *PEG10* frameshifting in the cytoplasm, by binding to mature mRNA and destabilizing the pseudoknot prior to ribosome engagement, or in the nucleus, by interacting with nascent transcripts and preventing proper pseudoknot formation during co-transcriptional folding.

To examine these possibilities, we compared the H3 TMO to a CapGap ASO design containing TMO-modified flanks and a central PS-DNA gap, which is potently RNase H-competent and induces target RNA cleavage. This architecture was intended to test whether nuclear RNase H activity contributes to the observed functional effects, consistent with the presence of RNase H in transcriptionally active nuclear compartments associated with RNA polymerase II. We also introduced pre-annealed ASO–RNA duplexes, which are known to localize predominantly in the cytoplasm (Hirano and Komatsu 2022) and would be expected to exchange with endogenous mRNA only if cytoplasmic strand displacement is a major contributor to ASO activity (Liu et al. 2025).

Functional analysis of PEG10 protein expression revealed that the CapGap design led to a significant reduction of gag and gag-pol levels, but no significant change in the gag-pol:gag ratio, consistent with RNase H–mediated degradation of *PEG10* mRNA and supporting a nuclear site of action (**Figure 5a-d**). In contrast, the fully modified H3 TMO selectively altered the gag-pol:gag ratio without inducing global PEG10 knockdown (**Figure 5a-d**), consistent with a non-degradative mechanism. Notably, when the H3 TMO ASO was delivered as a pre-annealed duplex with complementary RNA, its functional effects were partially abolished (**Figure 5d**), indicating that duplex delivery partially suppresses productive target engagement. These findings support a model in which ASOs interact with PEG10 mRNA in the nucleus, with a secondary cytoplasmic effect.

**Figure 5:**
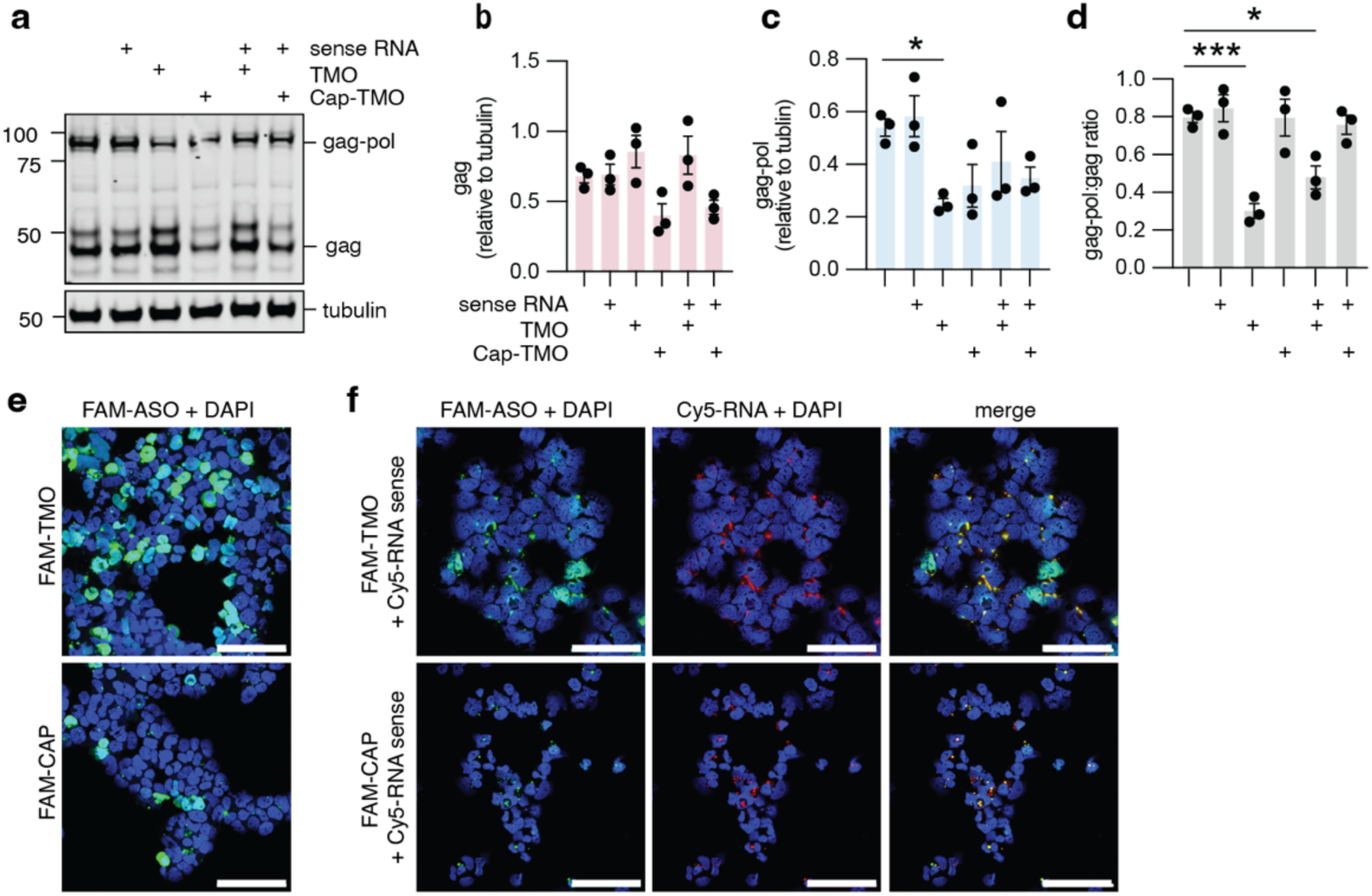
Duplexing of H3 TMO with RNA, which alters ASO localization, diminishes its effect on *PEG10* frameshifting. **a)** Representative western blot showing PEG10 levels with single or duplexed ASO chemistries. **b-d)** Quantification of gag (b), gag-pol (c), or the ratio of gag-pol:gag (d) for each condition in (a). For (a-d), experiments were performed three times, and statistics were determined by one-way ANOVA with multiple comparisons. **e-f)** Confocal microscopy of intracellular localization of fluorescently labeled oligonucleotides. HEK293 cells were transfected with labeled oligonucleotides and visualized 96 hours later following fixation and staining of nuclei with DAPI. Single-stranded H3 TMO or Cap-TMO ASO labeled with fluorescein (FAM) at the 5′ end (e), or RNA sense strand labeled with Cy5 at the 5′ end pre-annealed to either FAM-TMO or FAM-Cap-TMO (f) are shown. Images are representative of 3 experiments. Scale bar 100 μm.

To directly visualize intracellular localization, we performed confocal microscopy using fluorescently labeled oligonucleotides (**Supplementary Figure 3a**). First, fluorescently labeled ASOs were tested after transfection by western blot to ensure that they affected the gag-pol:gag protein ratio similarly to unlabeled ASO (**Supplementary Figure 3b-d, left**). FAM-labeled TMO and Cap-TMO were both predominantly nuclear (**Figure 5e**), whereas the Cy5-labeled RNA sense strand localized primarily to the cytoplasm, with enrichment in perinuclear regions consistent with endoplasmic reticulum association (**Supplementary Figure 4a**). In contrast, pre-annealed duplexes of TMOs displayed strong cytoplasmic co-localization of FAM and Cy5 signals (**Figure 5f**), consistent with impaired nuclear entry of duplex species, and showed a loss of effectiveness in their ability to influence the gag-pol:gag protein ratio (**Supplementary Figure 3b-d**). Of the additional chemistries tested, only PS-MOE and PS-LNA were capable of maintaining an effect on gag-pol:gag ratio when complexed with RNA (**Supplementary Figure 3a-d**). Together, these data support a model in which the leading H3 ASO engages PEG10 mRNA predominantly in the nucleus, but also in the cytoplasm.

### TMOs influence PEG10 production in neuronal cell lines

High levels of PEG10 gag-pol protein are observed in brain and spinal cord of murine models of familial ALS and in human spinal cords of ALS patients, while gag is not significantly changed (Whiteley et al. 2021; Black et al. 2023). Gag-pol is also specifically elevated in the neurodevelopmental disease Angelman’s syndrome (Pandya et al. 2021). Therefore, modulation of *PEG10* PRF could be an attractive therapeutic target in neurological disease. Because neurons may respond differently to ASO treatment when compared to traditional cell lines, SH-SY5Y cells were used as an approximation of a neuron-like cell line. Transfection of SH-SY5Y cells with the TMO H1 did not influence the ratio of gag-pol and gag protein abundance; however, TMO H3 had a modest effect on the ratio of gag-pol:gag proteins that was most evident at 200 nM (**Figure 6a-b**). Gymnotic delivery of these TMOs had no effect on the gag-pol:gag ratio (**Figure 6c-d**), indicating that in this cell-culture system, lipid transfection is necessary for the effect on *PEG10* translation.

**Figure 6:**
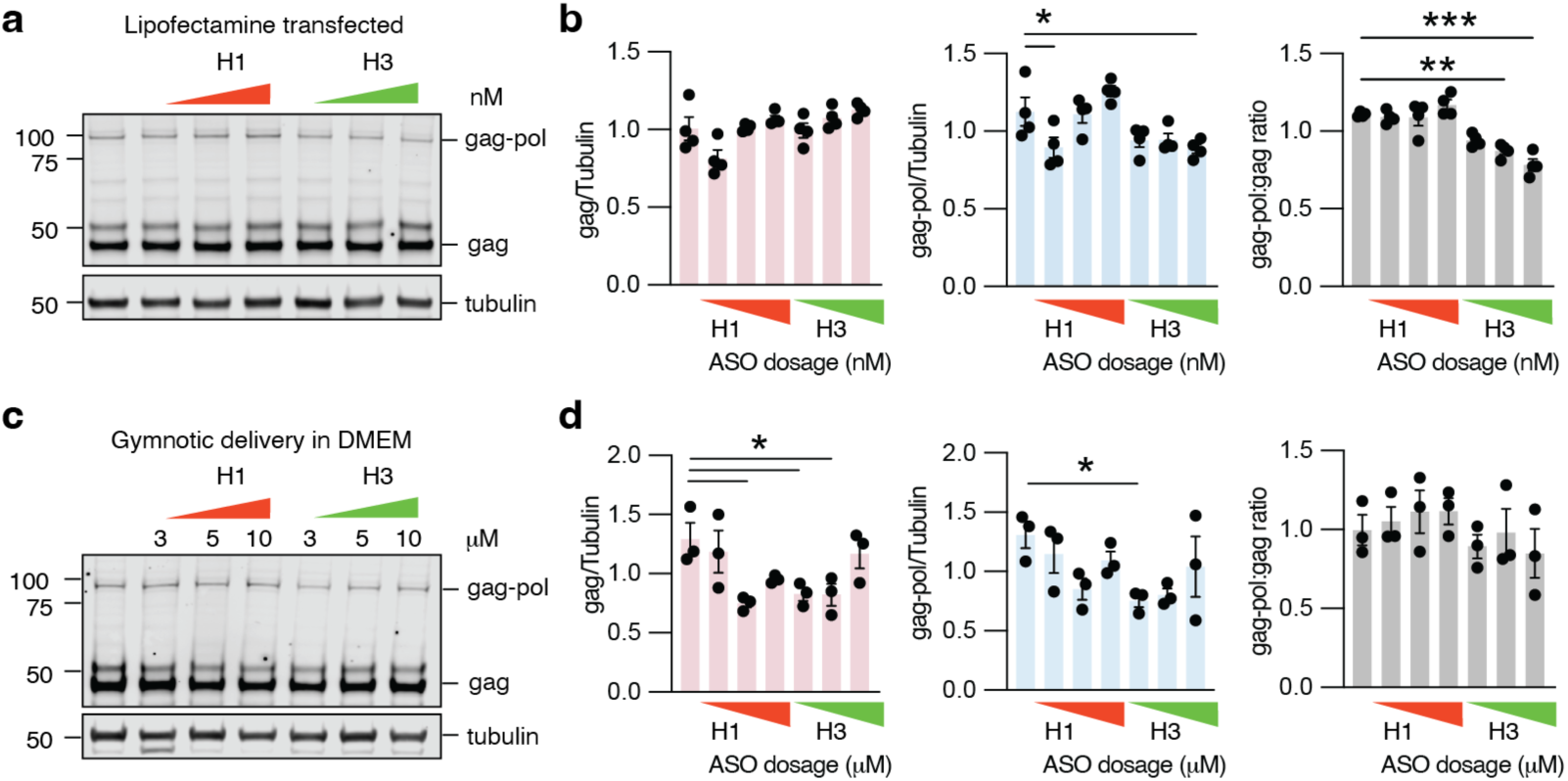
ASOs alter endogenous *PEG10* frameshifting in SH-SY5Y cells. **a)** Representative western blot of SH-SY5Y cells 72 hours after transfection with H1 and H3 TMO ASOs at either 0, 50, 100, or 200 nM concentration. Endogenous levels of PEG10 gag and gag-pol were probed. Shown is one of four representative blots. **b)** Quantitation of (left) gag, (middle) gag-pol, and (right) gag-pol:gag ratios from (a). Shown are mean ± SEM for four independent experiments. Significance was determined by one-way ANOVA with multiple comparisons. **c)** Gymnotic delivery of thiomorpholinos to SH-SY5Y cells. Cells were plated with the indicated concentration of ASO diluted in media for 7 days. Shown is one of 3 representative blots. **d)** Quantitation of (left) gag, (middle) gag-pol, (right) and the ratio of gag-pol:gag from (c) showing no major changes to PEG10 upon gymnotic delivery of ASOs. Shown are mean ± SEM for three independent experiments. Significance was determined by one-way ANOVA with multiple comparisons.

### Murine cells treated with frameshift inhibiting M3 ASO show a loss of gag-pol

Human and mouse PEG10 differ considerably at the amino acid level but retain the same predicted protein domain structure and mRNA pseudoknot (**Supplementary Figure 5a, Figure 7a**). However, a challenge of working with murine Peg10 is the inability of commercial antibodies to detect gag or gag-pol protein. Therefore, alternative strategies were used to assess the ability of TMOs to target murine Peg10. First, 3T3 cells were co-transfected with a plasmid encoding murine Peg10 containing an HA-tag on the N-terminus as well as murine-compatible M2, M3, or M4 TMOs (**Figure 7b**). While gag levels were largely unaffected by TMOs (**Supplementary Figure 5b**), gag-pol protein was dramatically decreased (**Supplementary Figure 5c**), leading to a lower apparent gag-pol to gag ratio (**Figure 7c**). To complement this strategy and assess the ability of ASOs to modulate endogenous murine *Peg10* translation, proteomics was performed on murine cells treated with ASO. Lewis Lung Carcinoma (LLC) cells were transfected with lipid, or M2, M3, or M4 ASO, then harvested for proteomics 48 hours later. To ensure Peg10 quantitation, a sample was included with Peg10 overexpression to trigger MS1 (Budnik et al. 2018; Yi et al. 2019). Five unique peptides were observed from murine Peg10: one from gag, and four from gag-pol (**Supplementary Figure 5a, black bars**). Only the M3 ASO showed an average gag-pol:gag peptide abundance ratio below 1.0, suggesting potency in preventing endogenous murine Peg10 from frameshifting (**Supplementary Figure 5d**). To extend our work into primary murine neurons, cortical neurons were isolated from E16.5 C57BL/6 mice and cultured. On day 4 of culture, cells were transfected with an AAV8 expressing HA-tagged murine Peg10; on day 5, cells were transfected with M3 TMO. On day 7, cells were harvested for western blot, which showed a dose-dependent decrease in the ratio of gag-pol:gag protein upon M3 transfection (**Figure 7d-e**). Together, these data suggest that pseudoknot-targeting TMOs represent an attractive means of modulating PEG10 frameshifting in neurons, which has therapeutic implications for diseases with particularly high levels of PEG10 gag-pol (Black et al. 2023; Pandya et al. 2021).

**Figure 7:**
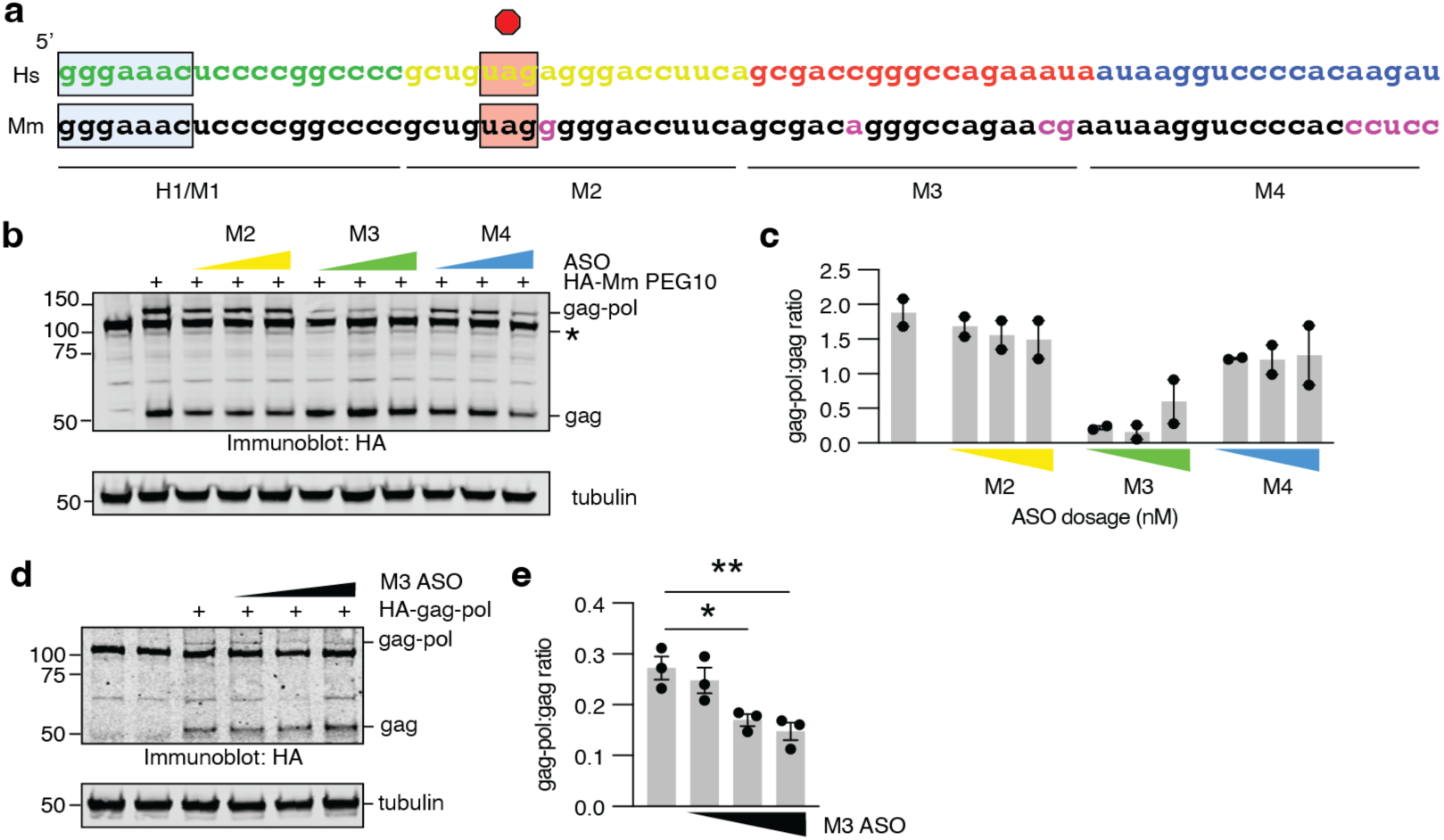
Murine *Peg10*-targeting ASOs similarly modulate frameshifting in cell lines and neurons. **a)** Schematic of murine *Peg10* pseudoknot sequence with differences between mouse and human highlighted in magenta below. Corresponding ASO sequences are shown below. **b)** Western blot of murine 3T3 cells simultaneously co-transfected with HA-tagged murine *Peg10* construct and either M2, M3, or M4 ASOs. Star denotes a nonspecific band. Peg10 was detected with HA-antibody. Shown is one of two representative experiments. **c)** Quantitation of results from (b). Gag and gag-pol levels are shown in Supplementary Figure 5. **d)** Western blot of murine cortical neurons derived from E16.5 embryos after infection with an HA-Peg10 expressing AAV8 followed by 48 hours of transfection with *Peg10*-targeting ASOs at 50, 100, or 200 nM. Peg10 was detected with HA-antibody. Shown is one of three representative experiments. **e)** Quantitation of gag-pol:gag ratio for results in (d). Gag and gag-pol levels are shown in Supplementary Figure 5. Statistics were determined by one-way ANOVA with multiple comparisons.

## Discussion

In this study, we establish *PEG10* -1 PRF as a tunable process that can be selectively modulated using antisense oligonucleotides designed to target the conformational dynamics of the stimulatory pseudoknot rather than by targeting the coding sequence itself. By integrating structure prediction, systematic antisense tiling, and multiple model systems, we identified a discrete vulnerability within the lower stem of the *PEG10* pseudoknot that governs frameshift efficiency. ASOs targeting this region robustly and selectively suppress gag-pol production with minimal impact on protein expression of gag, demonstrating that frameshifting can be functionally uncoupled from overall PEG10 expression.

A variety of ASO chemistries were capable of modulating *PEG10* frameshifting. While TMOs had the strongest effect on gag-pol:gag ratio, MOEs and LNAs also caused a significant reduction of *PEG10* frameshifting. Neither OMe or Cap-TMO ASOs significantly altered frameshifting. In the case of Cap-TMO, this could be consistent with degradation of the *PEG10* mRNA due to RNase H activation, resulting in a concurrent loss of both gag and gag-pol protein production; however, it remains unclear why OMe had so little effect. In conclusion, TMOs, MOEs, and LNAs would be an attractive starting point for further testing of frameshift modulating ASOs.

Mechanistic dissection using RNase H–active and –inactive ASO designs, pre-annealed duplexes, and fluorescence-based localization further suggests that productive ASO engagement with *PEG10* mRNA occurs in the nucleus, and to a lesser extent, in the cytoplasm. The persistence of ASO–RNA complexes through nuclear export and into the cytoplasm suggests that early structural perturbations can be propagated to the translational stage, ultimately biasing ribosomal decoding outcomes.

One remaining question concerns the ability of these ASOs to modulate gene expression in the absence of lipid-based transfection reagents. In SH-SY5Y cells, lipid-based transfection resulted in robust modulation of *PEG10* frameshifting, but incubation in media with micromolar concentrations of TMO were ineffective. While TMOs showed the strongest effect in multiple cell culture systems, they are relatively untested compared to alternative chemistries such as PS-MOE and PS-LNA. Future work is necessary to determine the feasibility of delivering TMOs in the absence of lipid transfection, which is a challenge shared amongst multiple ASO chemistries that are currently used in the clinic.

Given the selective elevation of PEG10 gag-pol in neurological disorders such as ALS and Angelman Syndrome (Black et al. 2023; Pandya et al. 2021), targeted modulation of *PEG10* frameshifting represents a promising and mechanistically precise therapeutic avenue. This may be particularly important for proteins like PEG10, where the frameshifted form of the protein has been associated with diseases, but the shorter form has also been found to be essential for reproduction (Shiura et al. 2025). More broadly, our work establishes a conceptual and experimental framework for regulating recoding events through antisense-mediated control of RNA structural dynamics, opening new opportunities to therapeutically target programmed ribosomal frameshifting in endogenous genes and viral genomes.

## Acknowledgements

A.M.W. and A.M.M. were supported by NIH R01 NS131660 and CDMRP AL240155. A.M.M. was supported by T32GM142607. M.M.S. and K.E.W. were also supported by CDMRP AL240155. We would also like to acknowledge the Shared Instruments Pool (SIP) at the Department of Biochemistry at CU Boulder (RRID: SCR_018986), the Biochemistry Cell Culture Core Facility (CCF) (RRID: SCR_018988), the Proteomics and Mass Spectrometry Core Facility (RRID: SCR_018992), and the BioFrontiers Institute’s Advanced Light Microscopy Core (ALMC) (RRID: SCR_018302). Proteomics was performed on a Thermo Q-Exactive HF-X supported by NIH S10-OD025267. Laser scanning confocal microscopy was performed on a Nikon A1R microscope supported by NIST-CU Cooperative Agreement award number 70NANB15H226 or the Nikon AXR Laser Scanning Confocal, which is supported by NIH Grant 1S10OD034320.We would like to thank Tom Cech for critical feedback on this manuscript. We would also like to thank Doug Dellinger, Ben Lunstad, and Logan Garner of Cirena for long RNA preparation guidelines.

## Author Contributions

Performed experiments: Kostov, Swanton, Waldon, Matthews, Ganguly, Ebmeier. Data analysis: Kostov, Swanton, Waldon, Matthews, Ganguly, Ebmeier, Whiteley, Caruthers. Prepared Materials: Kostov, Ciba, Schafer, Danielsen. Writing (original draft): Kostov, Swanton, Whiteley. Writing (editing): Kostov, Swanton, Matthews, Caruthers, Whiteley. Funding acquisition: Whiteley, Caruthers.

## Methods

### Oligonucleotide synthesis

All ASOs were synthesized on an ABI-394 synthesizer using Glen Research monomers for PS-DNA, PS-OMe, PS-MOE, and PS-LNA according to the manufacturer’s protocols. RNA sequences were prepared using TC-RNA chemistry (Dellinger et al. 2011), following the methodology developed by Cirena for long RNA synthesis (Dr. Oligo DNA RNA oligo synthesizer). All TMO sequences were prepared using a modified synthetic technology based on the previously published procedures (Langner et al. 2020), with modifications to all aspects of the synthesis cycle and downstream workup. To ensure suitability for safe application in living systems, the purified products were converted to their sodium salts using sodium acetate (0.3 M sodium acetate solution, BioUltra, for molecular biology) and subsequently purified from excess salts using an Amicon Ultra-15 15 mL centrifugal filter unit with a 3 kDa molecular weight cutoff. The final products were lyophilized from DNase-free water.

### Thermal Stability Study

Thermal denaturation experiments were performed using a Cary 100 Bio UV-VIS spectrophotometer, equipped with a 6×6 thermostatted multicell holder and a Peltier temperature controller. Oligonucleotides (ONs) and their complementary strands were mixed in equimolar ratios (1.0 µM per strand) in a buffer containing 100 mM NaCl, 50 mM NaH2PO4, and 1 mM EDTA at pH 7.2. The samples were transferred to 1 mL cuvettes and subjected to a heating cycle starting from 20°C and increasing to 100°C at a rate of 1°C per minute. After reaching 100°C, the samples were held at this temperature for 5 minutes before cooling down to 4°C at the same rate. Thermal denaturation measurements were taken after cooling. The denaturation curves were recorded at 260 nm, with a ramp rate of 0.5°C per minute. Data analysis and processing were carried out using Cary WinUV software. The melting temperature (Tm) values were determined by locating the peak in the first-derivative plots of absorbance versus temperature, with a precision of ±1°C.

### Cloning

Human PEG10 gag-pol (AA 1-708) containing an N-terminal fusion with mCherry and a C-terminal fusion with eGFP was generated from Gibson cloning (GeneArt HiFi Mastermix cat #A46629) into pCDNA3.1, then into the pCDNA5 plasmid (Life Technologies cat #V601020) for generation of stable cell lines. Murine HA-tagged Peg10 (AA 1-1006) for transient transfection into cell lines was generated in the pCDNA3.1 backbone through Gibson cloning. All generated constructs transformed into chemically competent DH5ɑ *E. coli* cells (Invitrogen). Transformed *E. coli* were plated on 100 μg/mL carbenicillin (Gold Biotechnology, cat #C-103–5) LB agar (Teknova, cat #L9115) plates overnight at 37 °C. Single colonies were picked and grown overnight in 5 mL LB Broth (Alfa Aesar, cat #AAJ75854A1) with carbenicillin at 37°C with shaking at 220 rpm. The following day, shaking cultures were mini-prepped (Zymo, cat #D4212) and sent for Sanger Sequencing (Azenta) or whole-plasmid sequencing (Plasmidsaurus and Azenta). Sequence-verified plasmids were then midi-prepped (Zymo, cat #D4201) for use in transfection.

Murine HA-tagged Peg10 (AA 1-1006) for AAV8-facilitated infection of murine cortical neurons was generated from pCDNA3.1 starting material and sent to Packgene for cloning and generation of AAV8 particles.

### Antibodies

Primary antibodies used for western blot included PEG10 (Proteintech rabbit polyclonal cat #14412-1-AP) used at 1:1000, Tubulin (Novus murine monoclonal DM1A cat #BB100-690) used at 1:10,000, and anti-HA tag (CST rabbit polyclonal cat #3724s) used at 1:1000. Secondary antibodies for western blot included anti-mouse IgG IRdye 680 (Licor goat polyclonal cat #926-68070) and anti-rabbit IgG IRdye 800 (Licor goat polyclonal cat #926-32211), both at concentrations of 1:20,000.

### Cell culture

Mammalian cells were maintained in the Biochemistry Cell Culture Facility at CU Boulder (RRID:SCR_018988). Cell lines were grown in complete media (DMEM (Gibco, cat #12800082), 10% FBS (Atlas Biologicals, cat #F-05000-DR), GlutaminePlus (Bio-techne Sales Corporation cat #R90210), and Penicillin-Streptomycin (Invitrogen, cat #15140163)). Once cells reached 100% confluency, cells were washed with PBS then treated with Trypsin-EDTA (0.05%) (Invitrogen, 25300120), until cells began to detach. Complete media was added to harvest the cells. Cells were then pelleted at 300 x g for 5 minutes and then the cell pellet was resuspended in fresh complete media. HEK293 cells were split at a 1:10 ratio for maintenance and SH-SY5Y cells were split at a 1:5 ratio for maintenance into new flasks. When plating for an experiment, 10 μL of final cell solution was combined with 10 μL of trypan blue (Gibco, cat #15250061) and live cells were counted via hemacytometer or using an automated cell counter (Corning, CytoSMART).

Stably expressing human PEG10 frameshift reporter cells were generated through the Flp-In system. In brief, Flp-In-293 cells (Invitrogen, cat #R75007) were cultured according to manufacturer instructions in complete media with zeocin (Gibco, cat #R25001). Cells were grown in a 6-well plate until they reached ∼70% confluency and zeocin media was removed and replaced with complete media. Cells were then transfected with 500 ng of pcDNA5/FRT plasmids or positive transfection control, pcDNA5/FRT chloramphenicol acetyltransferase (CAT) (Invitrogen, cat #V601020), in combination with 500 ng pOG44 (Invitrogen, cat #V600520) in Lipofectamine 2000 and Opti-Mem medium, according to manufacturer’s instructions. Once cells had grown to confluency, media was replaced with complete media with 100 μg/mL hygromycin B (Gibco, cat #10687010) to select for transfected cells. Cells were checked and media was replaced every 3-4 days until all untransfected cells had died.

For culture of murine cortical neurons, timed pregnancies were used to generate cells. Before embryo harvest, sterile cell-culture treated 12-well dishes were coated with 1 mL of mouse laminin (Invitrogen cat #23017-015) at 2.5 μg/mL final concentration and poly-D-lysine (Sigma cat #P7280) at 100 μg/mL final concentration diluted in PBS and filter sterilized, and kept in an incubator until use. E16.5 pregnant female C57BL/6 mice were humanely euthanized by CO2 and cervical dislocation and embryos taken out for neuron dissection and placed in ice-cold HBSS with 10mM HEPES. Brains were collected from embryos euthanized by rapid decapitation and cortices were peeled off of brains. Meninges were then removed and midbrain scraped off, along with hippocampus removal, via microscope-aided micro-dissection, all on ice in HBSS with HEPES. Isolated cortices were then chopped up into small pieces with scalpel and a maximum of three brains were combined into one tube containing fresh HBSS with HEPES. 500 μL of 2.5% stock solution of trypsin (Worthington cat #TRL3) was added and tube was incubated at 37°C for 30 minutes. After incubation, 500 μL of a 0.5% stock solution of DNAse (Sigma cat #DN25) was added and tube gently inverted to mix, then let sit at room temperature for 5 minutes. After incubation, media was removed and 5 mL fresh HBSS with HEPES was gently added to the tube to avoid breaking up brain chunks. Media was carefully changed twice more with HBSS and HEPES, then three times with HBSS with HEPES and 10% FBS, then three times with DMEM with 10% FBS and pen/strep, being careful each time to wait until brain chunks had settled. After equilibration into DMEM media, chunks were gently triturated while being careful not to introduce air bubbles using a fire-polished glass Pasteur pipet with bulb attached. After trituration, cells were pipetted through a 40 μm cell strainer placed at an angle in a 15 mL tube. 2 mL of DMEM with 10% FBS and pen/strep was added to the cells, and cells were then counted with hemacytometer.

7.5 x 10^5^ live cells were plated per well in 1 mL DMEM media with 10% FBS and pen/strep. Dishes were gently shaken up/down, then left/right to disperse cells. After one hour, cell attachment was confirmed by light microscopy and media was gently changed to NBActiv4 (BrainBits cat #NBActiv4500) supplemented with penicillin/streptomycin. Media was then changed every other day until harvest on day 7.

### Animals

C57BL/6 mice were obtained from Jackson Laboratories. Mice were housed at the University of Colorado, Boulder, according to Institutional Animal Care and Use Committee guidelines and in compliance with the Institute for Lab Animals’ guidelines for the humane care and use of laboratory animals. Timed pregnancies were set up between males and females between 8-35 weeks of age. All euthanasia was performed with CO2 and cervical dislocation according to Institutional Animal Care and Use Committee standards.

### Plasmid transfections and delivery of ASOs

Mammalian cells for PEG10 transfection were seeded in 12 or 6-well plates and grew until they reached ∼70% confluency. Cell media was changed to warmed transfection media containing DMEM (Gibco, cat #12800082), 10% FBS (Atlas Biologicals, cat #F-05000-DR), and GlutaminePlus (Bio-techne Sales Corporation cat #R90210). Cells were transfected with 1 μg (12-well plate) or 2.5 μg (6-well plate) plasmid DNA in Lipofectamine 2000 (Invitrogen, cat #11668027) and Opti-Mem medium (Invitrogen, cat #11058021), according to manufacturer’s instructions (see list of plasmids). 48 hours after transfection, cells were harvested for subsequent experiments.

For ASO transfection of cell lines, SH-SY5Y cells were seeded at 2.5 x 10^5^ cells per well of a 12-well plate. Cells were grown to 70% confluency for transfections. On the day of transfection, media was replaced with warmed transfection media (DMEM, 10% FBS, and 1X L-glutamine). Cells were then transfected with 50 nM, 100 nM, or 200 nM H1 or H3 ASO in Lipofectamine 2000 and Opti-Mem medium, according to manufacturer’s instructions. Cells were harvested for western blot analysis after 72 hours.

For infection of cortical neurons with AAV8, followed by ASO transfection, cortical neurons were first infected with 5 x 10^9^ CFU of virus per well on day 5, followed by a media change the next day. On day 6, cells were transfected with either 50 nM, 100 nM, or 200 nM M3 TMO using RNAiMax reagent (Life Technologies cat #13778100) according to manufacturer’s instructions.

### Western blot

Cells were collected and centrifuged at 300 x g for 5 minutes to harvest cell pellets. Cell pellets underwent two washes of PBS followed by centrifugation at 300 x g for 5 minutes each time. After washing, cell pellets were lysed in urea lysis buffer (8M urea (Fisher Chemical, cat #U153), 75 mM NaCl (Honeywell Fluka, cat #6003219), 50 mM HEPES (Millipore Sigma, cat #H3375) pH 8.5, 1x tab cOmplete Mini EDTA-free protease inhibitor cocktail tablet (Roche, cat #11836170001)). Lysed cells were briefly vortexed, centrifuged on a tabletop centrifuge and let sit at room temperature for 10 minutes then kept overnight at -20C°. Lysate was centrifuged for 10 min at 21,300 × g and the supernatant was collected.

Protein was quantified by BCA (Pierce, cat #23227). Western blot samples were prepared with 1x Laemmli sample buffer supplemented with βME (Sigma Aldrich, cat #M3148), and urea lysis buffer before SDS-PAGE. Samples were run in NuPage MES Running Buffer (Invitrogen, cat #NP000202, diluted to 1x in DI water) on a 4 to 12% NuPage Bis-Tris gel (Invitrogen, cat #NP0321). Following SDS-PAGE, the gel was wet transferred onto a nitrocellulose membrane (Amersham Protran, cat #10600009) for 45 minutes at 15V.

Membranes were blocked using LICOR blocking buffer (cat #927–70001) for 1 hour at room temperature. Membranes were incubated in primary antibody overnight at 4 °C in 1x TBST (50 mM Tris-Cl (MP Biomedicals, cat #MP04816100) pH 7.4, 150 mM NaCl, 0.1% Tween VWR, cat #M147-1L). Membranes were washed in 1x TBST 3 times in 5-minute intervals. Membranes were then incubated in LICOR secondary antibody in 1x TBST for 1 hour in the dark. Membranes were washed in 1x TBST 3 more times in 5-minute intervals, then the nitrocellulose membranes were visualized using LICOR Odyssey CLx. Data analysis was performed using LICOR ImageStudio Software.

### Fluorescence Microscopy

To study cellular uptake and localization of oligos, HEK293 cells were subjected to lipofection in above mentioned process in a 24 well plate. 72h post transfection, the cells were washed gently with 500 μL of DPBS twice. Next, 500 μL of Trypsin with EDTA was added to each well of the 24 well plate and the plate was placed in an incubator for 2-3 min. Once the cells were dislodged, 500 μL of DMEM media was added to each well to neutralize the solution. Then cells were collected and spun down at 1000 g for 10 min. After that, the supernatant was removed and 500 μL of fresh DMEM media was added. Next, cells were plated with ∼5000 cells/well seeding density in a 24 well SensoPlate (SensoPlate™ 24 well, PS, F-bottom, glass bottom, black, with lid, sterile, single packed, Greiner Bio-One, cat #82050-898) and incubated overnight for the cells to be attached on to the plate surface. The SensoPlates were used for this study as they are considered ideal for applications requiring low autofluorescence with exceptional optical clarity.

Next day, cells were fixed using 2% PFA solution by adding 400 μL in each well for 10 min at 37°C. Then the PFA solution was removed and the cells were washed 3 times with 1X PBS. Next, for permeabilization, 400 μL of 0.1% Triton X-100 (Invitrogen) in 1X PBS was added in each well and the cells were incubated at room temperature for 15 min. Again, cells were washed with 1X PBS three times and cell nuclei were stained with DAPI (Invitrogen). For DAPI staining, a 5 mg/mL DAPI stock solution was diluted to 300 nM in PBS. Approximately 400 µL of the diluted DAPI staining solution was added to each well and the cells were kept incubated for 15 minutes with the lid on at RT. Then the cells were again washed with 1X PBS three times.

Next, cells were imaged in confocal microscopy (Nikon AXR) using 405/488 and 561 laser lines.

### Mass spectrometry

Murine LLC cells were transfected with TMO and lysed 48 hr later in 8M urea buffer with protease inhibitor. In one sample, lysate was mixed in a 95:5 ratio of untransfected LLC lysate to LLC lysate transfected with murine HA-tagged Peg10 construct as a spike-in control. Approximately 100–200 μg of each sample was aliquoted and delivered to the Proteomics and Mass Spectrometry Core Facility in the Department of Biochemistry at the University of Colorado, Boulder, for TMT labeling. Protein samples were reduced and alkylated with the addition of 5% (w/v) sodium dodecyl sulfate (SDS), 1% (w/v) sodium deoxycholate, 10 mM tris(2-carboxyethylphosphine) (TCEP), 40 mM 2-chloroacetamide, 50 mM Tris-HCl, pH 8.5 and incubated shaking at 1000 rpm at room temperature for 60 min, then cleared via centrifugation at 17,000 × g for 10 min at 25 °C. Lysates were digested using the SP3 method (Hughes et al. 2014). Briefly, 400 μg carboxylate-functionalized speedbeads (Cytiva Life Sciences) were added to approximately 100 μg protein lysate. Addition of acetonitrile to 80% (v/v) induced binding to the beads, then the beads were washed twice with 80% (v/v) ethanol and twice with 100% acetonitrile. Proteins were digested in 50 mM Tris-HCl buffer, pH 8.5, with 1 μg Lys-C/ Trypsin (Promega) and incubated at 37 °C overnight. Tryptic peptides were desalted using HLB Oasis 1 cc (10 mg) cartridges (Waters) according to the manufacturer’s instructions and dried in a speedvac vacuum centrifuge. Approximately 12 μg of the tryptic peptide from each sample was labeled with TMT-Pro 16-plex (Thermo Scientific) reagents according to the manufacturer’s instructions. The multiplexed sample was cleaned up with an HLB Oasis 1 cc (30 mg) cartridge. Approximately 50 μg multiplexed peptides were fractionated with high pH reversed-phase C18 UPLC using a 0.5 mm × 200 mm custom packed Reprosil Pur C18 1.9 μm 120 Å (Dr. Maisch) column with mobile phases 0.1% (w/v) aqueous ammonia, pH 10 in water and acetonitrile (ACN). Peptides were gradient eluted at 20 μL/min from 2 to 50% ACN in 50 min concatenating for 12 fractions using a Waters M-class UPLC (Waters). Peptide fractions were then dried in a speedvac vacuum centrifuge and stored at –20 °C until analysis.

### Mass spectrometry analysis

High pH peptide fractions were suspended in 3% (v/v) ACN, 0.1% (v/v) trifluoroacetic acid (TFA) and approximately 1 μg tryptic peptides were directly injected onto a reversed-phase C18 1.7 μm, 130 Å, 75 mm × 250 mm M-class column (Waters), using an Ultimate 3000 nanoUPLC (Thermos Scientific). Peptides were eluted at 300 nL/min with a gradient from 4 to 25% ACN over 120 min then to 40% ACN in 5 min and detected using a Q-Exactive HF-X mass spectrometer (Thermo Scientific). Precursor mass spectra (MS1) were acquired at a resolution of 120,000 from 350 to 1500 m/z with an automatic gain control (AGC) target of 3E6 and a maximum injection time of 50 milliseconds. Precursor peptide ion isolation width for MS2 fragment scans was 0.7 m/z with a 0.2 m/z offset, and the top 15 most intense ions were sequenced. All MS2 spectra were acquired at a resolution of 45,000 with higher energy collision dissociation (HCD) at 32% normalized collision energy. An AGC target of 1E5 and 120 milliseconds maximum injection time was used. Dynamic exclusion was set for 20 s with a mass tolerance of ±10 ppm. Raw files were searched against the Uniprot Mouse database UP000000589 downloaded June 4, 2021 using MaxQuant v.2.0.3.0. Cysteine carbamidomethylation was considered a fixed modification, while methionine oxidation and protein N-terminal acetylation were searched as variable modifications. All peptide and protein identifications were thresholded at a 1% false discovery rate (FDR). Peptide abundances for each sample’s identified Peg10 peptides, including those that only differed in charge but had the same peptide sequence, were first compared to the average of that peptide for all samples, in order to control for idiosyncratic abundances of different Peg10 peptides. Each peptide was also assigned to gag or pol regions of Peg10 dependent on location. Then, those averages were used to generate a pol:gag ratio for each sample.

---

**Supplementary Figure 1:**
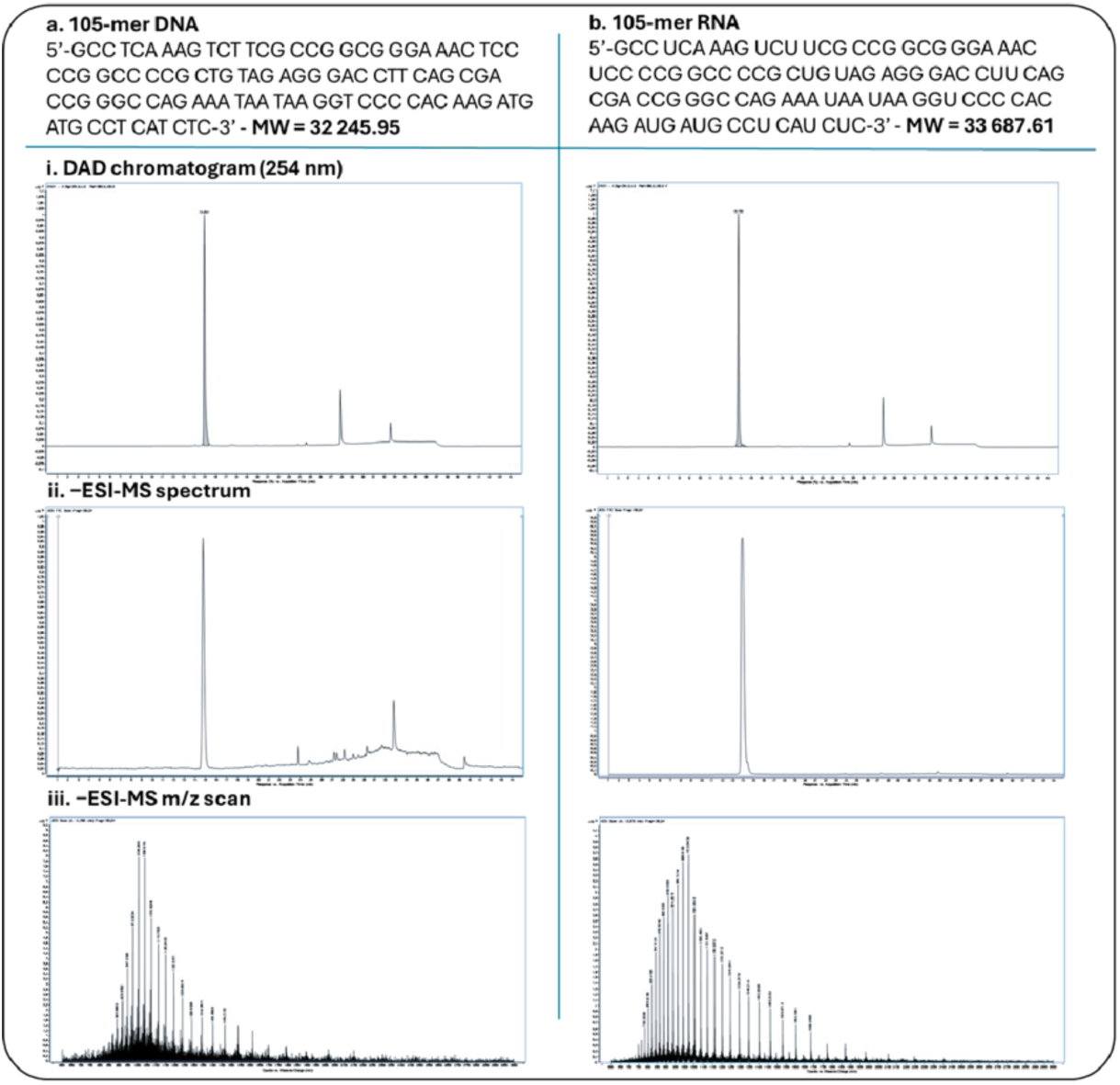
Analytical characterization of the 105-nt synthetic model sequence mimicking the *PEG10* pseudoknot. **(a)** Synthetic DNA version of the 105-nt model sequence. **(i)** Diode-array detector (DAD) chromatogram recorded at 254 nm. **(ii)** Negative-mode electrospray ionization mass spectrum (−ESI-MS). **(iii)** Deconvoluted −ESI-MS spectrum confirming the expected molecular mass. **(b)** Synthetic RNA version of the 105-nt model sequence. **(i)** DAD chromatogram at 254 nm. **(ii)** −ESI-MS spectrum. **(iii)** −ESI-MS m/z scan.

**Supplementary Figure 2:**
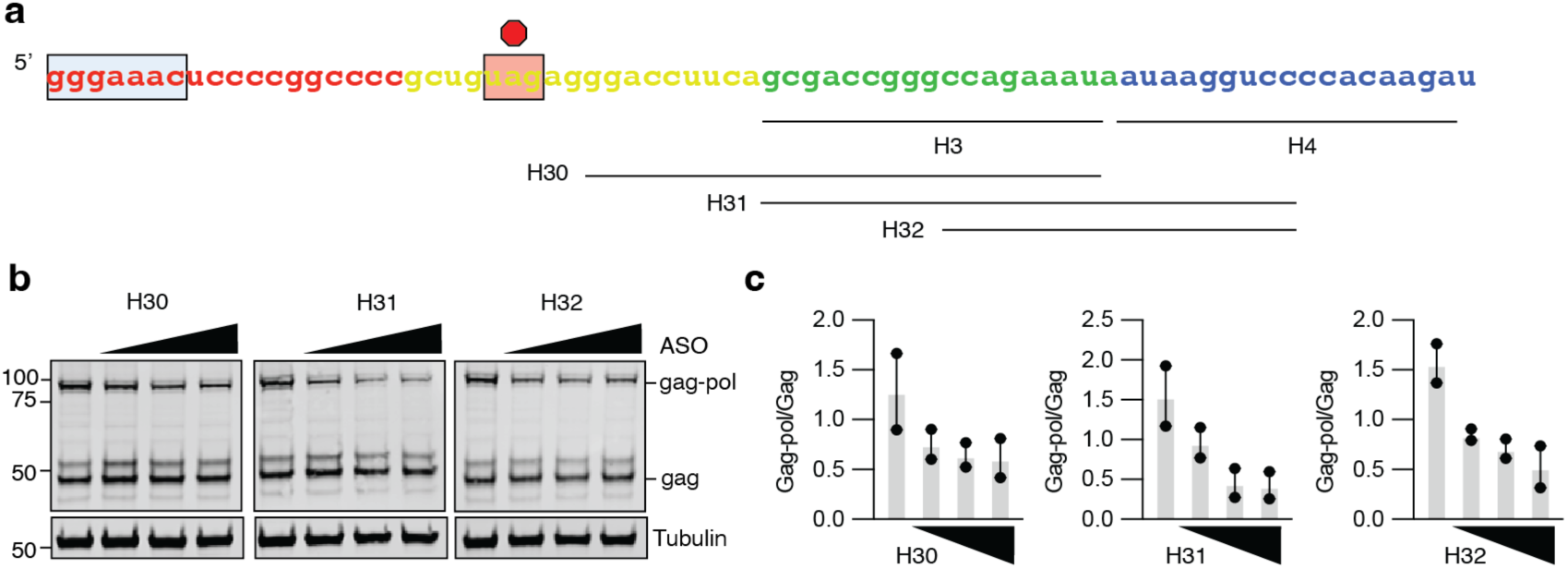
Second-generation thiomorpholino ASOs are not meaningfully better than first-generation ASOs in modulating gag-pol:gag ratio. **a)** Schematic of second-generation human *PEG10*-targeting ASOs as compared to H3 and H4. H30 and H31 are longer ASOs; H32 is an 18-mer that spans regions of H3 and H4. **b)** Representative western blot comparing efficacy of second-generation ASOs to modulate PEG10 frameshifting. HEK293 cells were transfected with 0, 50, 100, or 200 nM ASO as in Figure 3 and harvested 72 hours later for western blot. Shown is one of two representative blots. **c)** Quantitation of the gag-pol:gag ratio for each of the three second-generation ASOs. Data are from two representative experiments.

**Supplementary Figure 3:**
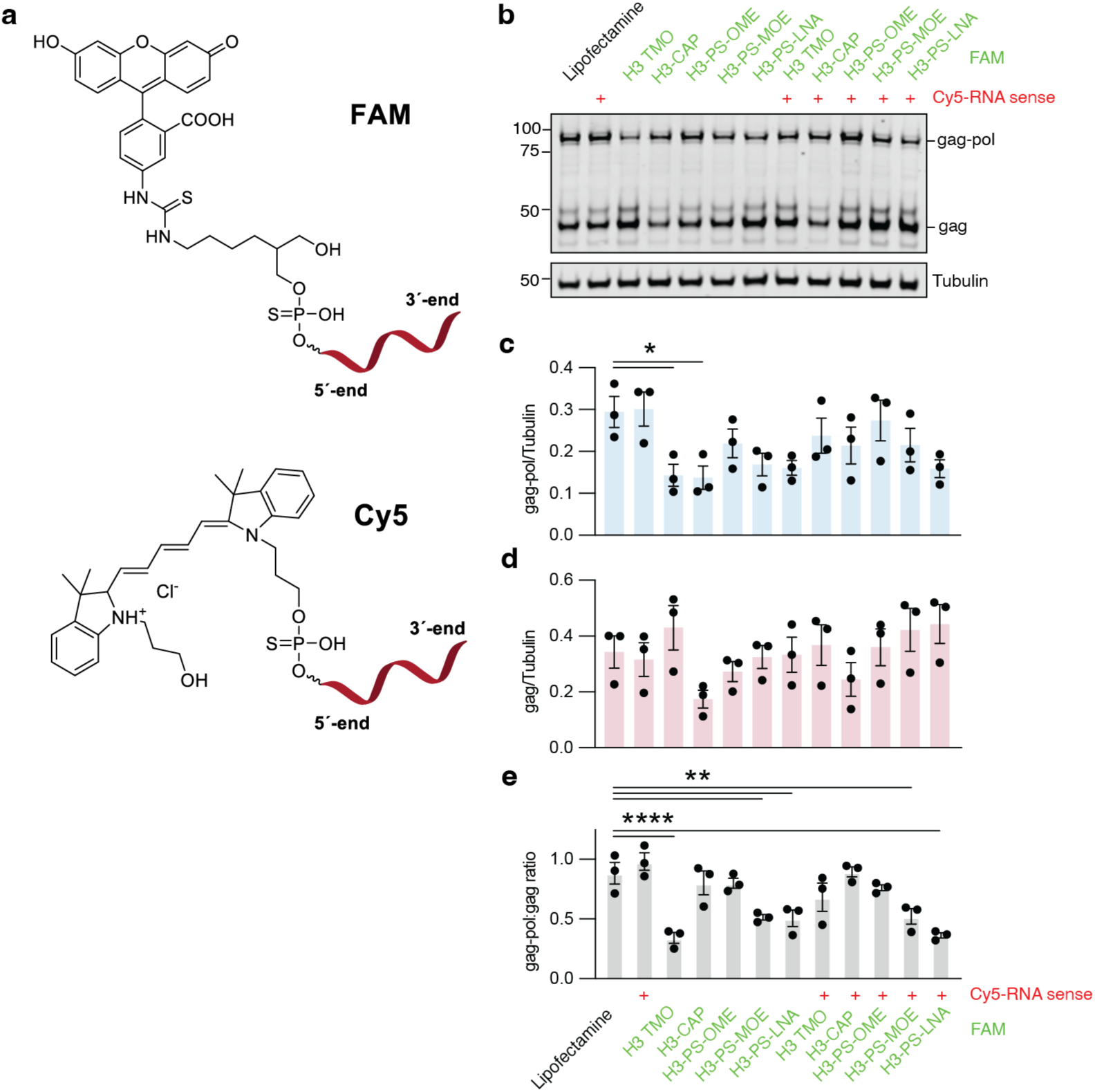
Modification of ASOs with fluorophores results in similar effects on PEG10 frameshifting, but formation of RNA:ASO duplexes mitigates their effect. **a)** Schematic of fluorophore connection to ASO and RNA showing thiophosphate linkage between fluorophore and 5’ end of oligonucleotide. **b)** Example western blot with FAM-labeled ASOs with and without pre-annealing to Cy5-labeled sense sequence of RNA 72 hours after transfection into HEK293 cells at 100 nM. Shown is one of three representative blots. **c-e)** Quantitation of gag-pol (c), gag (d), and the ratio of gag-pol:gag abundance (e) from (b). Statistics were determined by one-way ANOVA with multiple comparisons.

**Supplementary Figure 4:**
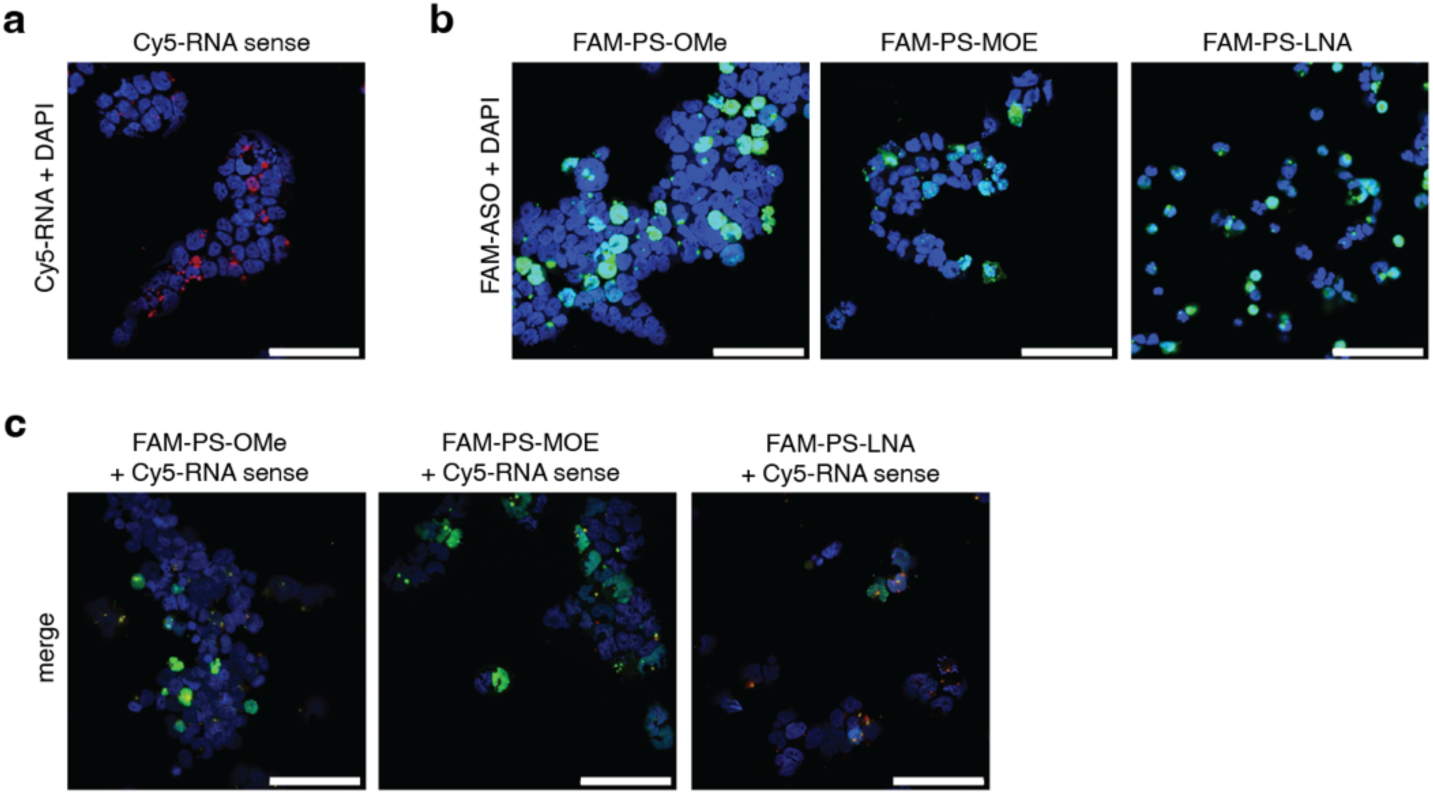
Pre-annealing of OMe, MOE, and LNA ASOs with RNA changes their localization. **a)** Micrograph of HEK293 cells transfected with Cy5-labeled RNA. **b)** Micrograph of HEK293 cells transfected with FAM-labeled PS-OMe, PS-MOE, or PS-LNA. **c)** Micrograph of HEK293 cells transfected with pre-annealed mixtures of Cy5-labeled RNA with either FAM-labeled PS-OMe, PS-MOE, or PS-LNA. For (a-c), scale bar is 100 μm and images are representative of 3 experiments. Within each sub-panel, microscope and collection parameters were kept the same between ASO types.

**Supplementary Figure 5:**
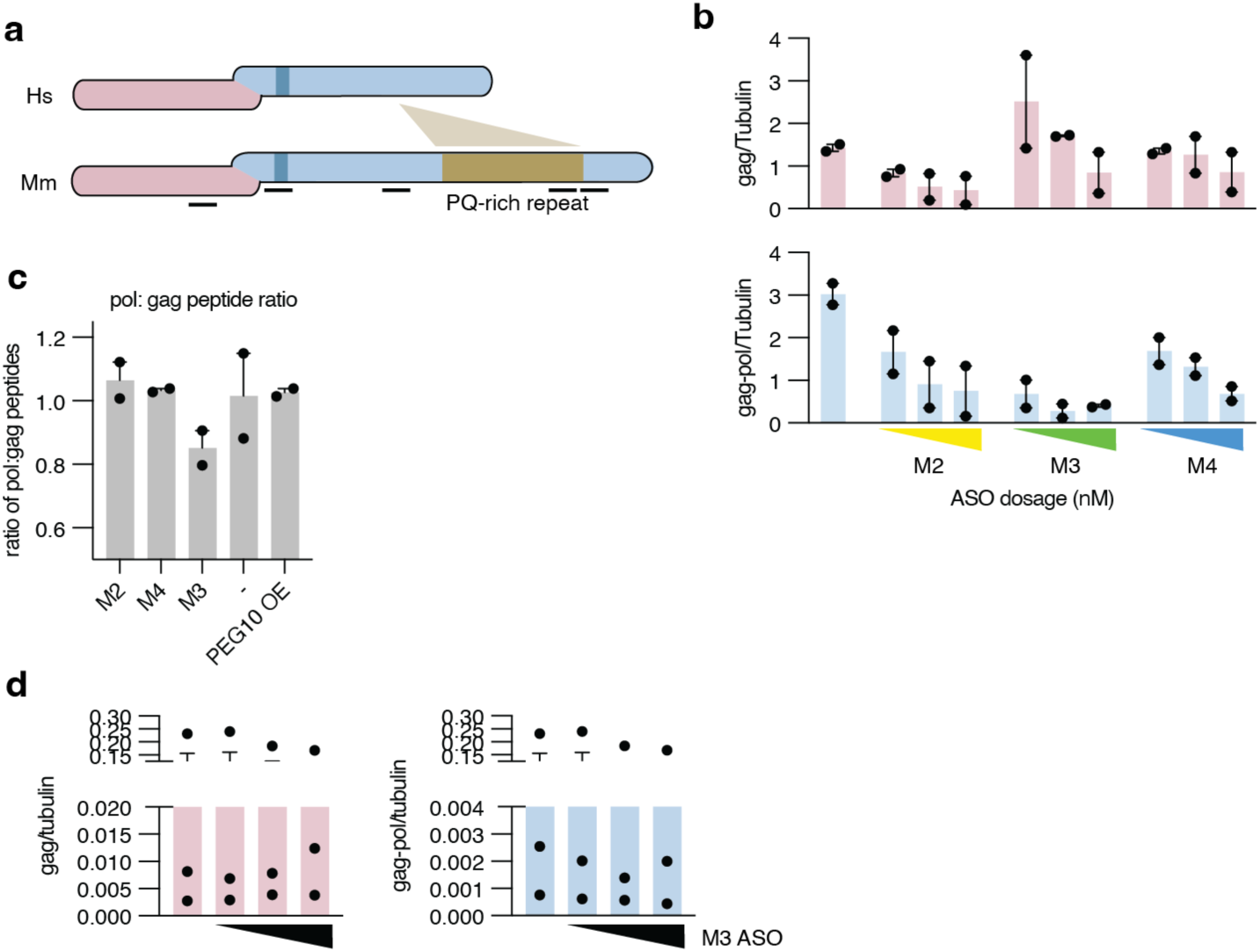
Murine *Peg10* is similarly affected by Loop 2-targeting TMOs. **a)** Schematic showing the general amino acid sequence divergence between PEG10 in human and Peg10 in mouse. In the middle of the pol region of murine Peg10 is a proline and glutamine-rich (PQ-rich) repeat region. Black bars at the bottom represent the approximate locations of peptides that were identified by mass spectrometry of LLC cells in (d). **b)** Gag (top) and gag-pol (bottom) levels quantified for transfected 3T3 cells shown in Figure 7c. Statistics were performed via one-way ANOVA with multiple comparisons. **c)** Murine LLC cells were transfected with 100 nM ASO and harvested 48 hours later for mass spectrometry analysis of endogenous murine Peg10 protein. To trigger MS2 quantitation of Peg10, a sample was included with 5% lysate from cells overexpressing (OE) murine HA-tagged Peg10 protein. Peptide abundances for each sample’s identified Peg10 peptides were first compared to the average of all samples. Each peptide was assigned to gag or pol regions. Then, those averages were used to generate a pol:gag ratio for each sample. Data are from two independent transfections. **d)** Gag (top) and gag-pol (bottom) levels quantified for transfected murine cortical neurons shown in Figure 7d. Statistics were performed via one-way ANOVA with multiple comparisons. Gag and pol levels are shown with two-piece y-axis because of variability of Peg10 signal across experiments.

